# *intSDM* : an *R* package for building a reproducible workflow for the field of integrated species distribution models

**DOI:** 10.1101/2022.09.15.507996

**Authors:** Philip Mostert, Ragnhild Bjørkås, Angeline J.H.M. Bruls, Wouter Koch, Ellen C. Martin, Sam W. Perrin

**Affiliations:** Department of Mathematical sciences, Norwegian University of Science and Technology, 7491, Trondheim, Norway; Centre for Biodiversity Dynamics, Norwegian University of Science and Technology, 7491, Trondheim, Norway; Norwegian Biodiversity Information Centre, Trondheim, Norway; Gjærevollsenteret, Norwegian University of Science and Technology, 7491, Trondheim, Norway

**Keywords:** Citizen science, Data integration, Integrated species distribution model, Reproducible workflow, Spatial point process

## Abstract

1. There has been an exponential increase in quantity and type of biodiversity data in recent years, including presence-absence, counts, and presence-only citizen science data. Species Distribution Models (SDMs) have typically been used in ecology to estimate current and future ranges of species, and are a common tool used when making conservation prioritisation decisions. However integration of these data in a model-based framework is needed to address many of the current large-scale threats to biodiversity.
2. Current SDM practice typically underutilizes the large amount of publicly available biodiversity data and does not follow a set of standard best practices. Integrating different data types with open-source tools and reproducible workflows saves time, increases collaboration opportunities, and increases the power of data inference in SDMs.
3. We aim to address this issue by (1) proposing methods and (2) generating a reproducible workflow to integrate different available data types to increase the power of SDMs. We provide the *R* package *intSDM*, as well as guidance on how to accommodate users’ diverse needs and ecological questions with different data types available on the Global Biodiversity Information Facility (GBIF), the largest biodiversity data aggregator in the world.
4. Finally, we provide a case study of the application of our proposed reproducible workflow by creating SDMs for vascular plants in Norway, integrating presence-only and presence-absence species occurrence data and climate data.

## 2 Introduction

There has been an unprecedented increase in biodiversity information over the last decade as a result of the exponential rise in digital technology to collect and store data effectively (Michener & Jones, 2012). Analyzing this massive collection of data has the potential to help us in our efforts to answer a multitude of ecological questions on a far larger scale than could previously be achieved.

A major difficulty in employing these data at large however lies in the heterogeneity of the individual datasets collected, making combining them into a single-modelling framework, notably species distribution models (SDMs) challenging (Kelling *et al*., 2009; König *et al*., 2019; Simmonds *et al*., 2020). Individual datasets are often collected with a specific purpose in mind, leading to different spatio-temporal resolutions, sampling protocols and variable names between each dataset. A substantial amount of available data is collected by citizen scientists, which is often opportunistic in nature and known to contain a multitude of different biases (Sicacha-Parada *et al*., 2021). As a result, significant care needs to be taken when combing these datasets; neglecting the biases in analysis of them could lead to poor inference (Simmonds *et al*., 2020).

Integrated species distribution models (ISDMs) have been proposed as a solution to the problem of data heterogeneity: allowing researchers to combine data from disparate sources under a single modelling framework by assuming that each is derived from a common underlying distribution (Heberling *et al*., 2021; Isaac *et al*., 2020). A multitude of benefits have been found from implementing these models. Initially, they were used to show that the effect of biases typical in opportunistic occurrence records may be reduced by both including covariates or flexible spatial terms to account for the sampling biases, when analyzing them in conjunction with structured survey data (Dorazio, 2014; Simmonds *et al*., 2020). Furthermore, these models expand the scope or scale of interest, increase the precision and accuracy of models, improve inference about potential distribution shifts of organisms in space and time and allow data use to extend beyond original intent at the time of collection, blending different sampling methods and dataset properties (Dorazio, 2014; Fithian *et al*., 2015; Miller *et al*., 2019; Simmonds *et al*., 2020).

Typical ISDM analyses are tailored towards small-scale studies, and only use a subset of all data available. Despite this, the tools and computational resources needed to analyze big data using data-integration methods on a large scale are certainly accessible and available today (Farley *et al*., 2018). We therefore provide the tools and recommendations for developing a workflow required to estimate large-scale ISDMs in an automated and reproducible framework. The implementation of our proposed workflow is available through the *R* package (R Core Team, 2023) *intSDM*, which is designed to combine structured and unstructured data available on GBIF (Global Biodiversity Information Facility, the largest biodiversity data aggregator in the world).

Procedures and steps required to build reproducible workflows and pipelines to transform raw data to obtain periodic estimates of species occupancy have recently been described by both Boyd *et al*. (2023) and Cervantes *et al*. (2023). We follow the structure of their workflows in a similar fashion, but put emphasis on the methods regarding using multiple disparate datasets, and estimating large-scale integrated species distribution models.

Herein we report a case study of the application of our reproducible workflow to create species distribution models of vascular plants in Norway by integrating two sources of information: presence-only and presence-absence data using the *intSDM R* package. A detailed analysis of the case study is also provided as a vignette within the *R* package.

## 3 Methods

### 3.1 *intSDM* R package

Our proposed workflow is presented through the *R* package, *intSDM*. The package is designed to construct ISDMs at large by following the conceptual workflow steps described as follows:

1. Obtain heterogeneous occurrence data and covariate layers.
2. Process, clean and filter data.
3. Estimate an integrated model for the data.
4. Preform model assessment and selection.
5. Create summaries and communicate results.

This subsection is dedicated to describing the functions of the package, as well as the considerations practitioners should take when developing these types of workflows. An overview of the toolkit available within the *intSDM* R package is provided in Figure 1.

**Figure 1.**
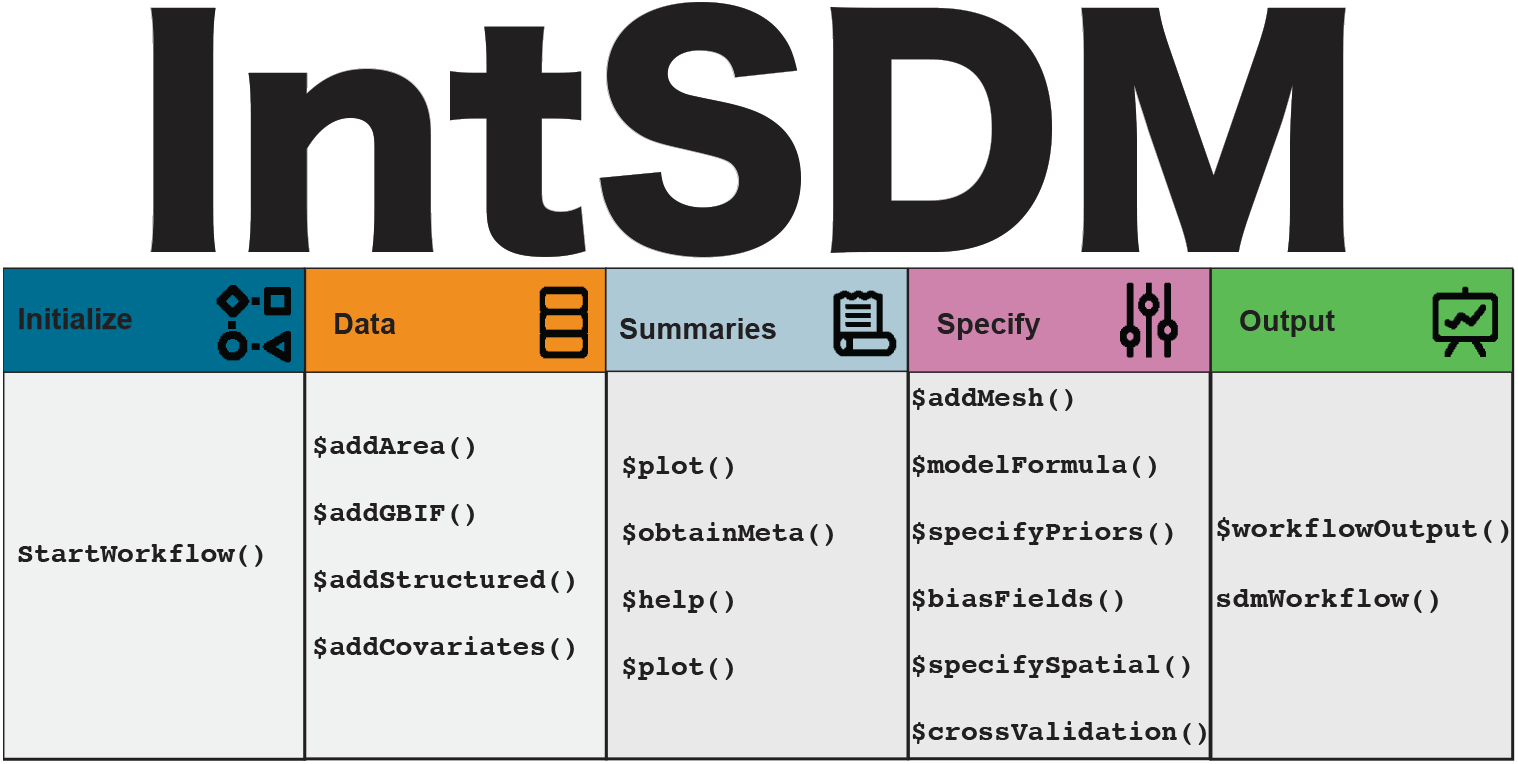
An overview of the functions used in the R package intSDM. These functions are categorized by one of five steps related to construction of the workflow: initialization, data, summaries, options or output.

The package is currently available on both CRAN and github, and may therefore be downloaded using either on of the following scripts:

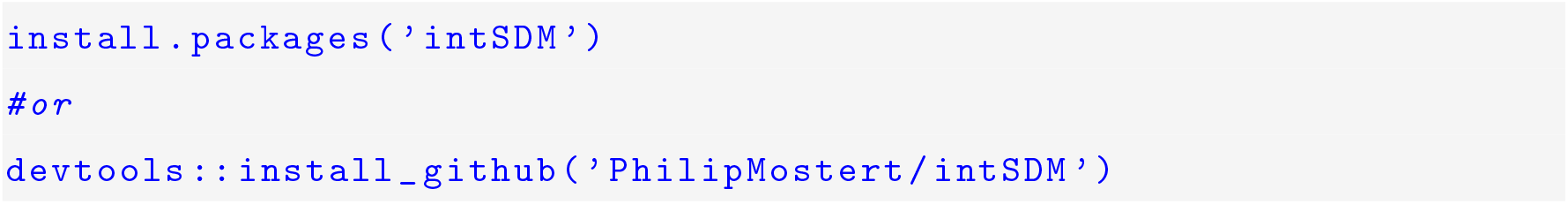

### 3.2 Functionality of package

The first function considered in the package is `startWorkflow`, which initialises the workflow. The arguments of this function are designed to specify non-model components of the workflow, such as the names of the species considered in the workflow and the study domain, as well as other project options such as directories and project names.

The default for this function is to create estimates for each species independently, however setting the argument `Richness = TRUE` estimates a multi-species model. This allows the user to obtain maps of estimated of species richness across the study area. An example of an analysis estimating species richness using the *intSDM R* package is presented as a vignette within the package.

The output from this function is an *R6* (Chang, 2021) object containing different slot functions (which are represented by the ‘`.$functionName’` notation) used to customize the different components of the workflow, breaking it down into manageable chunks. These slot functions are illustrated in figure 1, and are related to different parts of the workflow. Documentation and help related to these these slot functions is easily obtained by calling `.$help`.

Once all the parts of the workflow have been specified, the outputs of the workflow may be obtained using the function, `sdmWorkflow`. This function will return either a list of model outputs, or save it to a pre-defined directory, depending on the `Save` argument in `startWorkflow`.

#### 3.2.1 Identify and process data

Data are fundamental for our workflow, and two types of data are necessary to estimate species’ distributions: geolocated observation data detailing the sampling locations for the species across some geographic domain, and environmental covariate layers describing the environmental conditions at each location across the same domain.

Observation data may be obtained from numerous different online repositories such as *eBird* (Sullivan *et al*., 2014) or *iNaturalist* (Nugent, 2018). Within our setting, we focus on GBIF (GBIF, 2023) as it serves as an aggregated entry point for these and many other data sources.

This repository hosts a massive amount of data derived by a multitude of different data collectors and institutions using a variety of different sampling protocols in their collection process – although the data are mostly opportunistically collected citizen science records (Amano *et al*., 2016). Even though these observations are known to contain biases, their use in research is increasing and they remain a vital source of information to supply into our SDMs (Feldman *et al*., 2021).

Data from GBIF may be added to our workflow using the `.$addGBIF` function. The function’s arguments are related to specifying the observation process of the dataset (one of presence-only, presence-absence or counts), as well as additional arguments to filter and process the data from GBIF using tools from the *rgbif* (Chamberlain *et al*., 2022) package (see `?rgbif::occ_data` for the filtering options available).

A primary objective in this step is separating the different occurrence datasets into different observation processes based on the sampling protocol considered in the collection protocol. This information is often known and can be located within the datasets’ metadata file (Isaac & Pocock, 2015). The observation process for a given dataset is specified in the workflow using the function’s *datasetType* argument.

However, the most important part of using GBIF-obtained data in a large-scale workflow, is that it is FAIR (findable, accessible, interoperable, and reusable) (Wilkinson *et al*., 2016), and follows data standards set by Biodiversity Information Standards (International Working Group on Taxonomic Databases, 2006), such as Darwin Core, Ecological Metadata Language, and the Biological Collection Access Service standards (Heberling *et al*., 2021), making it easy to obtain and standardize in an automated procedure.

Data sharing of raw data on large online repositories is a key ingredient for data integration, and has become more prevalent across ecology over the last decade (Culina *et al*., 2020). Despite this, not all data are readily available for open-research (König *et al*., 2019; Michener & Jones, 2012). As a result, researchers may need to obtain data from secondary locations. Sometimes occurrence data may be obtained from other sources such as country- or institution specific databases or through collaboration efforts (Michener & Jones, 2012). These data would typically be high-quality survey data, and thus a valuable asset in conjunction with the citizen science records if the researcher is able to obtain them.

Non-*GBIF* data may be added to the workflow using `.$addStructured`. The use of the function is similar to that of `.$addGBIF`, with arguments related to specifying the observation process for the data, as well as specifying the response and variable names within each. However, the downside is that these data require more effort to bring into the workflow given that they may not be standardized in the same way that *GBIF* data is, and pipelines to transfer them directly to the workflow may not exist.

In addition to observation data, this step of the workflow includes compiling environmental, habitat, or other variable data to be included in the models. They are typically collected via remote sensing or geographic information system (GIS) methods; possible sources of such data include WorldClim (Fick & Hijmans, 2017), Copernicus (Copernicus, 2023), CHELSA (Karger *et al*., 2017) and Light Detection and Ranging (LiDAR). These variables need to be selected with care prior to any analysis, being considerate of both the goals of the model and the underlying biology of the species studied (Araújo & Peterson, 2012).

WorldClim environmental layers may be added to the workflow using the `.$addCovariates` function. The arguments here are used to specify the name of the covariate, resolution and function to apply to the data. Like the species occurrence data, environmental data other than WorldClim may be preferable. These environmental layers may be added using the function’s *Object* argument.

#### 3.2.2 Data documentation

Discovered data needs to be documented in order for it to be reproducible. This becomes a critical step as data are combined for use in analysis and terms of use and quality control vary. Adequate data documentation is a key component of methods reproducibility (Zurell *et al*., 2020), and metadata standards have been proposed to facilitate tracking information on data authorship, preparation (i.e., cleaning, changing, or validating), and model estimation (Merow *et al*., 2019). Given the wide range of species’ observational data and environmental data available for inclusion in SDMs, properly documenting the sources, metadata, and scales are necessary for reproducibility. Documenting data discovery and inclusion is also necessary to assess the comparative value of SDMs when different datasets are used to model the distribution of a same species.

Obtaining metadata from the workflow may be done using the `.$obtainMeta` function. This function provides the citations for the WorldClim covariates and each of the GBIF obtained datasets used in the analysis.

#### 3.2.3 Customization of the model

The model structure of the ISDM may be customized with some of the slot functions. The functions `.$specifySpatial` and `.$specifyPriors` allow the user to specify the priors for the spatial effects, and the fixed and random effects respectively. Strong priors for the effects in these types of models are often needed to reduce overfitting and improve overall model fit.

The function `.$modelFormula` allows for two arguments, *modelFormula* and *biasFormula* to assist in specifying the formula for the covariates associated with the process-level model and the covariates associated with the presence-only observation models used to account for bias respectively.

#### 3.2.4 Estimation of the Integrated Species Distribution Model

Applied SDM use is now well established, and there are a multitude of different methods to estimate these models (see for example, Fletcher and Fortin (2018)). Given the novelty of the field of integrated species distribution models, bespoke *R* packages and software to estimate the models appropriately are limited. Implementation of these models in the past has typically been done by assuming a point process framework, and by estimating a joint-likelihood through either maximum likelihood estimation (Dorazio, 2014; Fithian *et al*., 2015; Koshkina *et al*., 2017) or through Bayesian models using Markov chain Monte Carlo (MCMC) techniques (Miller *et al*., 2019; Zulian *et al*., 2021). However there are a variety of different methods to estimate ISDMs, for example: combining results from independent models, treating the results from one model as a covariate in another, using one dataset to derive an informed prior for another in a Bayesian setting, or by simply pooling the datasets together (Fletcher Jr *et al*., 2019).

The *PointedSDMs R* package (Mostert & O’Hara, 2023) has been proposed to estimate these complex models in an easy-to-use framework. *PointedSDMs* is built around the *R-INLA* package (Martins *et al*., 2013), which uses the integrated nested Laplace approximation (INLA) methodology (Rue *et al*., 2009) to approximate Bayesian latent Gaussian models. INLA is preferred to traditional Bayesian approximation techniques for continuous space models, for its speed and computational efficiency in estimating Gaussian random fields via the stochastic partial differential equation (SPDE) approach (Lindgren *et al*., 2011), which are used to account for unmeasured covariates and potential autocorrelations. *intSDM* uses *inlabru* (Bachl *et al*., 2019), an *R* package which builds wrapper functions around *R-INLA* to assist in the construction and estimation of spatial point process models.

Our integrated model is defined as in Isaac *et al*. (2020), where the hierarchical state-space model is the combination of a process model, used to reflect the “true” distribution of the species, with separate observation models for each of the datasets, where the likelihood for each of these models is dependent on the underlying sampling protocol of the corresponding dataset. Here, we treat the species’ “true” distribution as an inhomogeneous Poisson process, with an intensity function given by *λ* (*s*) = *e*^*η*(*s*)^, which describes the expected number of species at some point *s*. The log of this intensity is given by:

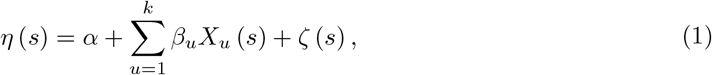

where: *α* is an intercept term, *X*_*u*_ (*s*) are spatially varying environmental variables with corresponding coefficients *β*_*u*_, and *ζ* (*s*) is a spatially varying Gaussian random field used to account for unmeasured covariates and potential spatial autocorrelation in the model. This linear predictor may be extended by adding variables to reflect biases or additional random fields. As a result, the expected number of species across some area Ω is therefore given by:

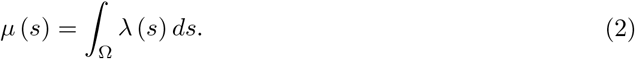

The observation processes for our state-space model depend on the type of data at hand, which typically come in the form of count data, presence-absence or presence-only data. Following Isaac *et al*. (2020) we model our count data in this framework as a Poisson point process with expected value given by the integral of the intensity function over the study area, and the presence-absence data with a Binomial likelihood with a *cloglog* link function, which allows us to relate the probability of occurrence of the species to the log of the intensity function. Given that citizen science data are often collected opportunistically and may contain a multitude of different sampling biases (Sicacha-Parada *et al*., 2021), we model the presence-only data from a thinned Poisson process to account for imperfect detection in the collection process.

#### 3.2.5 Validation and assessment of models

Validation of statistical models using independent data is vital in assessing model performance. However, model assessment and selection for ISDMs is incredibly difficult to complete since methods such as cross-validation and variable selection may not perform well under this framework (Zipkin *et al*., 2021).

Nevertheless, validation of results and model comparison is an important step in any scientific analysis. Therefore our workflow incorporates two types of cross-validation: a spatial-block cross-validation (using functionality from the *blockCV* R package [Valavi *et al*., 2019]) as well as a leave-one-one cross-validation (see for example [Mostert & O’Hara, 2023]). Cross-validation for the workflow may be specified by using `.$crossValidation`.

#### 3.2.6 Model summaries and outputs

The outputs of the ISDMs give us an understanding of how the underlying environment drives species richness and distribution across an area, and are typically applied to some form of biodiversity assessment (such as selecting areas for restoration and protection).

These results can then be used to make future projections on which species are most sensitive to what environmental conditions, and how the changes in these conditions could force range shifts across time. Such knowledge could be useful e.g., for management practitioners when identifying important areas for protection of biodiversity in the future, or when assessing risks of biological invasions (Guisan *et al*., 2013).

A popular output from these types of models are prediction maps of the intensity function of the model, which gives a reflection of the occurrence rate or the intensity of the species across a map. A useful way to extend the outputs of these models is by adding a temporal component to the model could reflect how this abundance changes over a time period.

Outputs from our model may be chosen using the `.$workflowOutput` function, which allows the user to specify different outputs from the workflow. These include obtaining the *R-INLA* model, predictions of the intensity function of the model, maps of the intensity function and cross-validation results.

### 3.3 Motivating case study

As a case study, we used our reproducible workflow for integrating disparate datasets to create a species distribution map for vascular plant species in Norway (this particular workflow is illustrated in figure 2). Such a map could be used for several purposes, for instance when locating new areas to protect for conservation. According to the Norwegian Red List of 2021, 22% of vascular plant species in Norway are considered threatened, i.e., placed in the Red List categories Critically Endangered (CE), Endangered (EN) or Vulnerable (VU) (Norwegian Biodiversity Information Centre, 2021). A major goal when designating new protected areas, both in Norway (Norwegian Biodiversity Information Centre, 2021) and globally (International Union for Conservation of Nature, 2022) is to protect species diversity and prevent extinction. It is essential to be able to accurately map the distribution of threatened species to locate areas of importance. To do this, all available data in various standards and qualities must be combined in an effective way.

**Figure 2.**
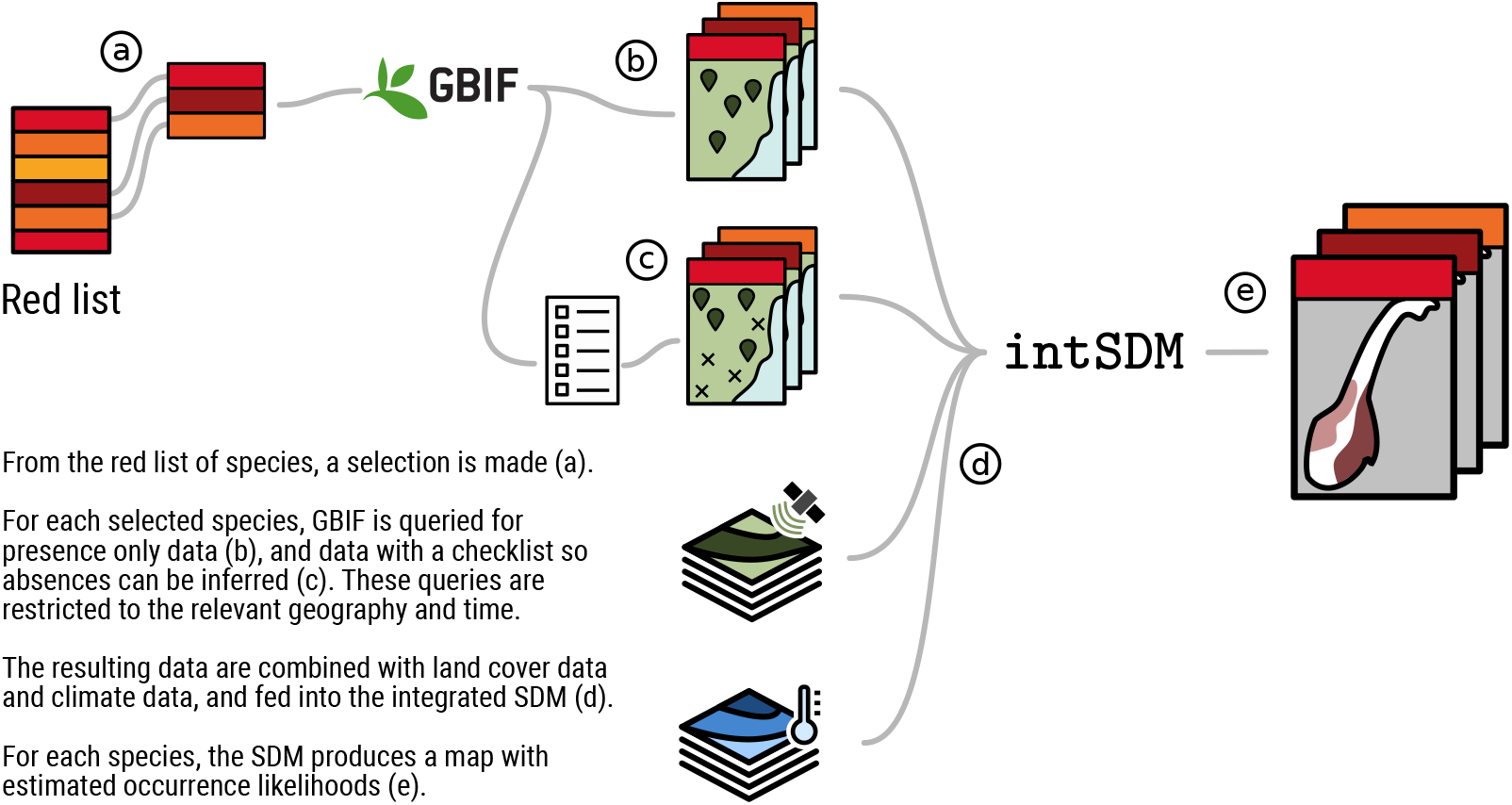
Visual summary of the workflow considered for our case study. The steps begin with a selection of species (a), a GBIF query of presence-only data (b) and presence-absence data (c), and a combination of covariate data (d). The package intSDM is used in the final step to estimate occurrence likelihoods (e).

For this example, we obtained presence-only data published by the Norwegian Biodiversity center through GBIF (The Norwegian Biodiversity Information Centre, 2023). We selected the three most abundant species in the data also categorised as being threatened on the Norwegian Red List (Norwegian Biodiversity Information Centre, 2021); *Arnica montana L*. (VU), *Fraxinus excelsior L*. (VU), and *Ulmus glabra Huds*. (EN). We further filtered any observations with a coordinate uncertainty greater than 50 m. Since these data contain no absence records, and are prone to high sampling biases, we assume these data are the outcome of a thinned Poisson process from an underlying log-Gaussian Cox Process (LGCP). We combined the species occurrence data with one environmental variable (annual mean temperature) in the ISDM to produce a map with estimated relative abundance across Norway for each species.

In addition, we supplemented the model by including two additional presence-absence datasets (modelled as Bernoulli random variables), also obtained from GBIF, which were collected and provided by *NTNU University Museum* (Norwegian University of Science and Technology, 2023) and *University of Oslo* (University of Oslo, 2023). These data used standardised cross-lists containing most vascular plants in Norway which, in addition to providing information about presences, also allowed for inference of species absences.

### 3.4 Implementation of case study

In this section we provide a brief example of how the *intSDM* package is used to implement our case study. We first begin the workflow by using the `startWorkflow` function. This requires us to specify the coordinate reference system considered, as well as the scientific names of the three species. After that, we use the `.$addArea` function to specify the study domain of the analysis.

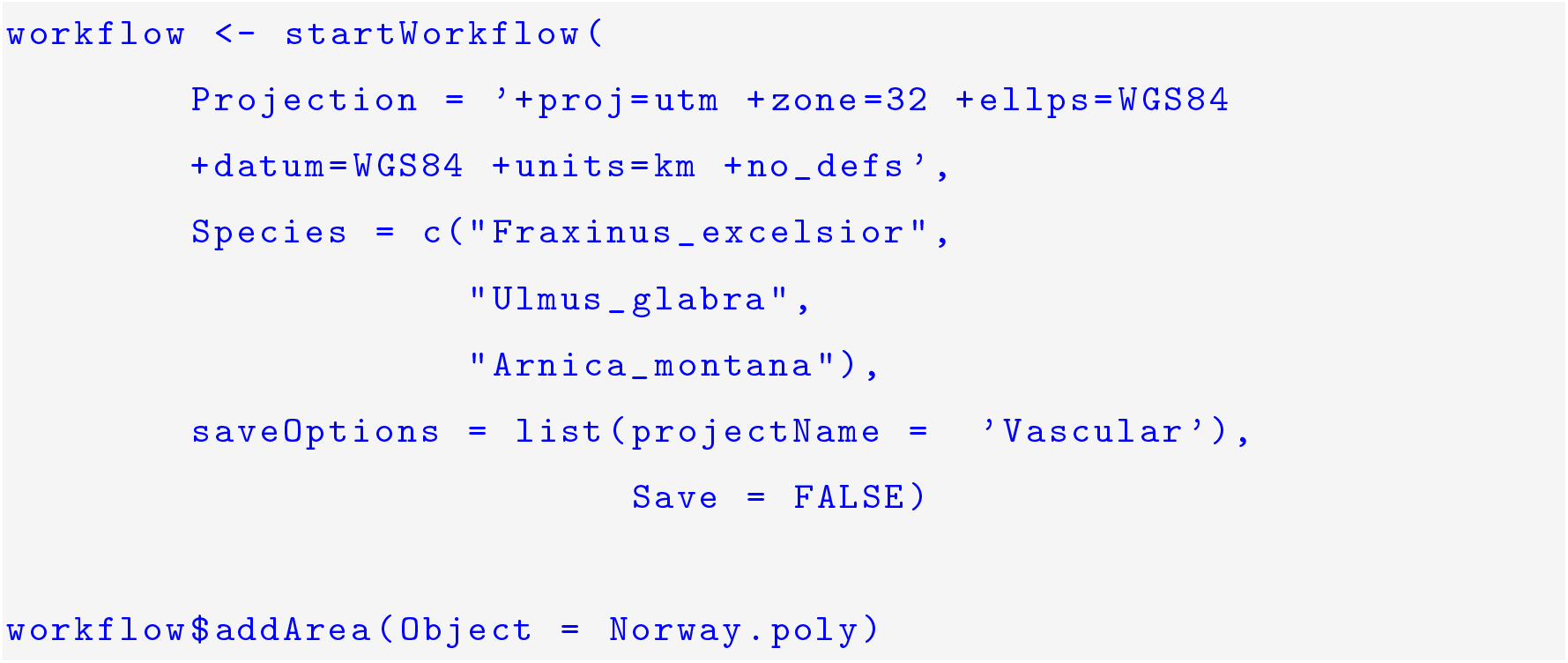

The species occurrence data may be added using the `.$addGBIF` function. The ellipse argument (…) links to rgbif’s `occ_data`, which allow us to filter and process the data. In this example, we specify arguments for *limit, coordinateUncertaintyInMeters* and *datasetKey*.

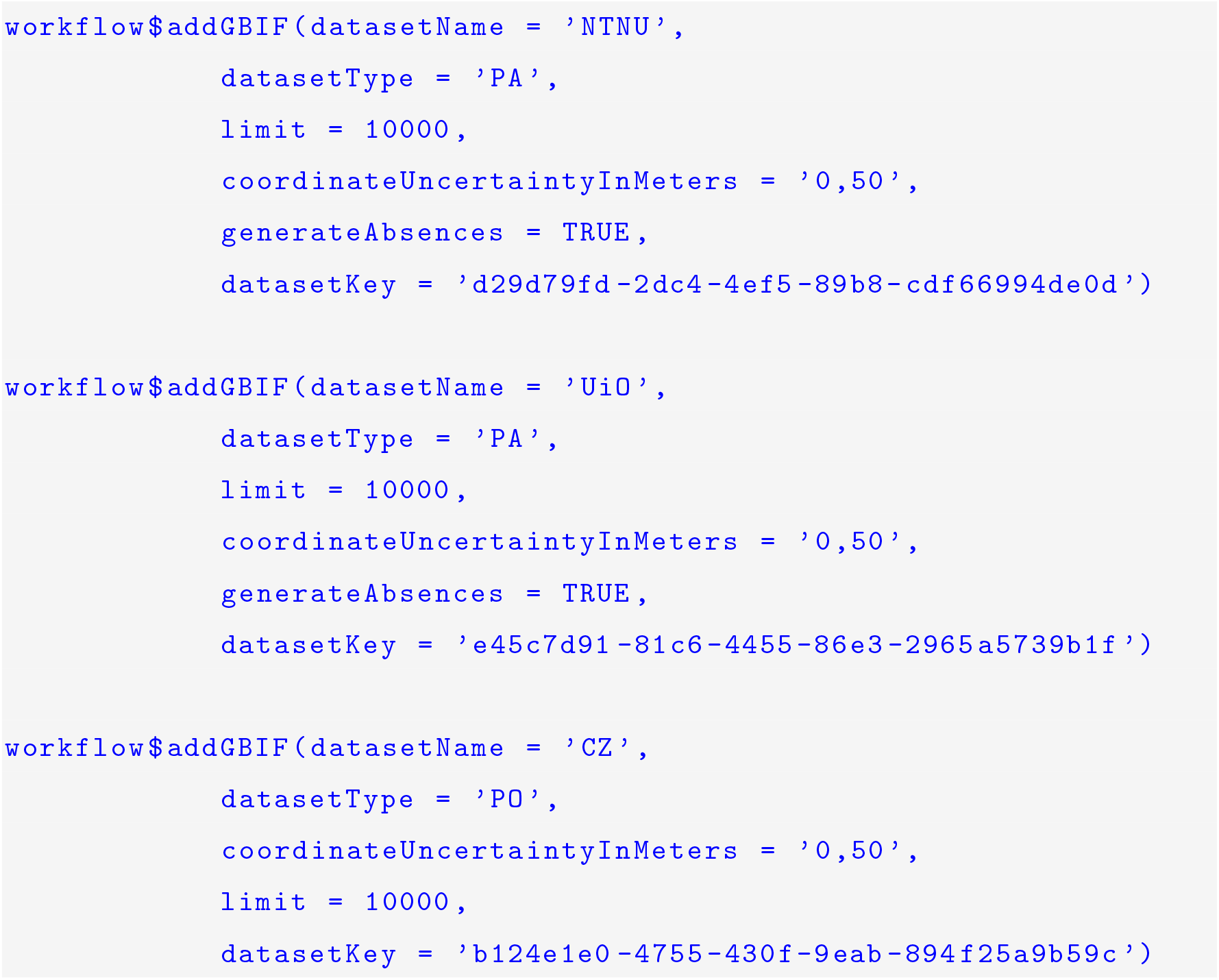

WorldClim covariates are added using the `.$addCovariates` function. For this case case study we selected the average temperature across Norway at a 5 minutes of a degree resolution, and standardized the raster data.

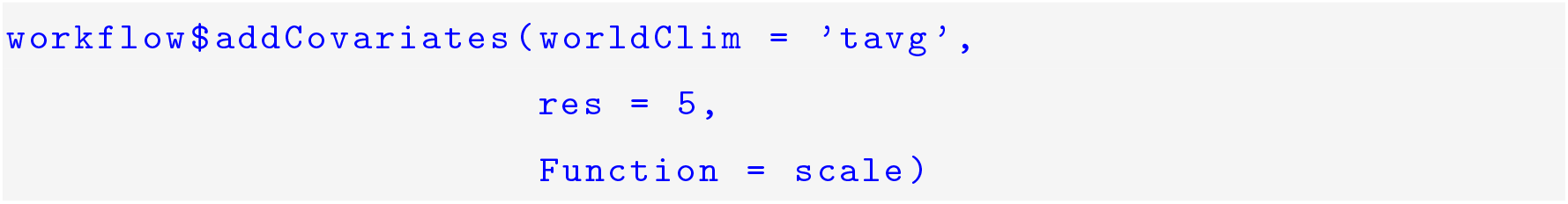

Our ISDM included a shared Gaussian random field between the datasets to account for potential spatial autocorrelation. In order to estimate this, we are required to create a mesh object, which can be done using the `.$addMesh` function. We used penalizing complexity priors for the range and sigma parameters for the Matérn covariance function of the random field (Simpson *et al*., 2017), which are designed to reduce over-fitting in the model by penalizing the model towards the base model. By doing this, we assumed that the probability of the range parameter of the covariance function being less than 100km was 10%, and the probability that the sigma parameter was greater than 1 was 10%. This was specified using the `.$specifySpatial` function. We also included a second random effect for the citizen science data using the `.$biasFields` function to reflect spatial-biases in the sampling protocol, as suggested in Simmonds *et al*. (2020). The same penalizing complexity priors were used for this spatial effect as we used for the shared spatial effect.

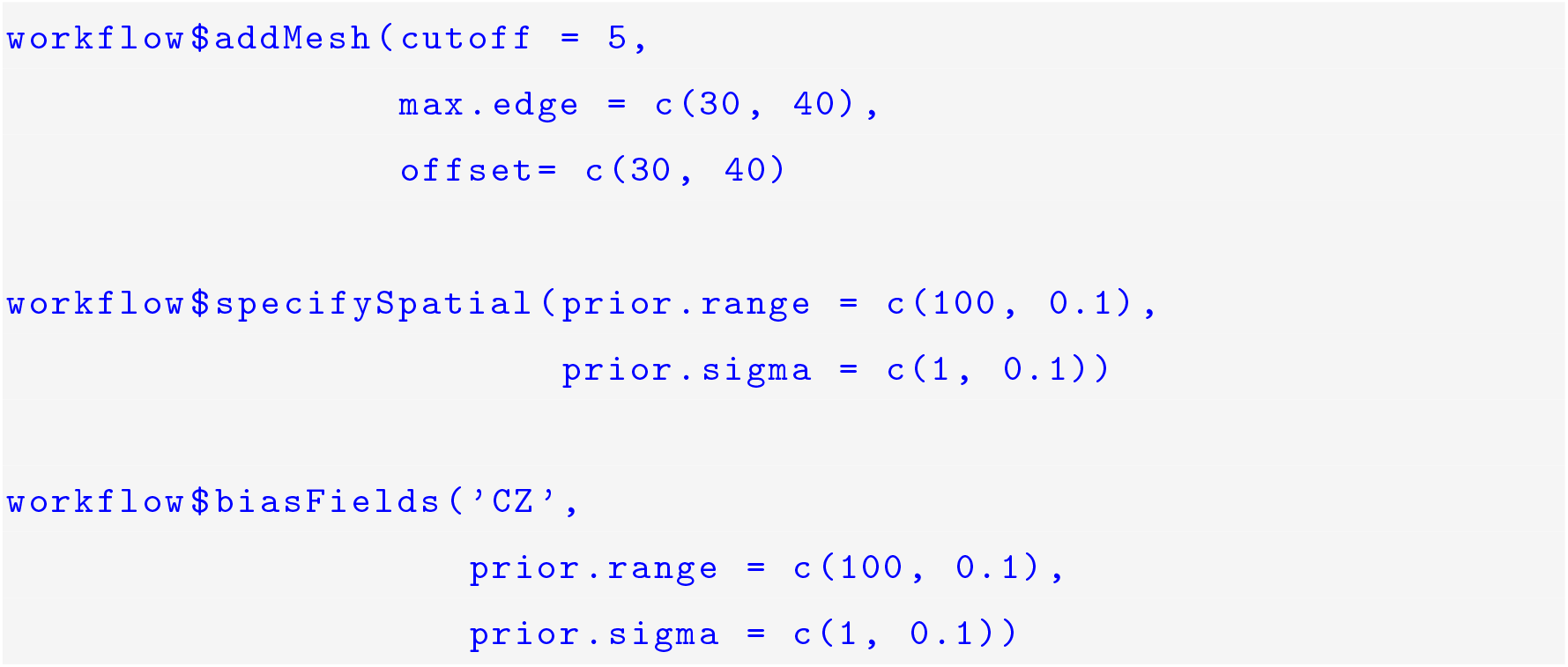

The output from this script were distribution maps reflecting the relative abundance for the three studied species and the bias fields, which was specified using the `.$workflowOutput` function. Additional *R-INLA* options are then added using the `.$modelOptions` function.

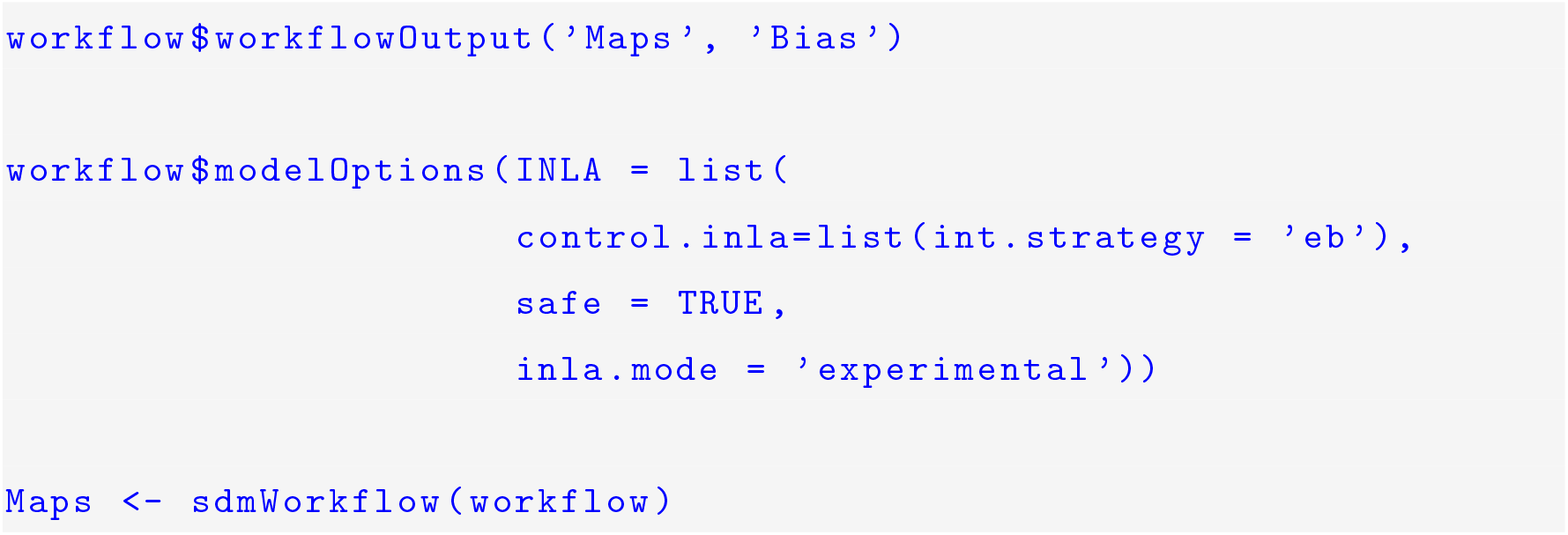

Finally the workflow was implemented using the `sdmWorkflow function`, which produced the maps as illustrated by figure 3. The predictions suggest similar distribution patterns for *Fraxinus excelsior* and *Ulmus glabra*, where they have a greater intensity along soouthern Norway and along the coast, but a low intensity in the Northern parts of Norway. The predicted intensity for *Arnica montana* suggests that it is more likely to occur in only the southern parts of the Country. The predicted bias fields for the three fields all look very similar, with a higher mean towards the southern part of Norway, and a lower mean towards the Northern section where there were little presence-only records collected.

**Figure 3.**
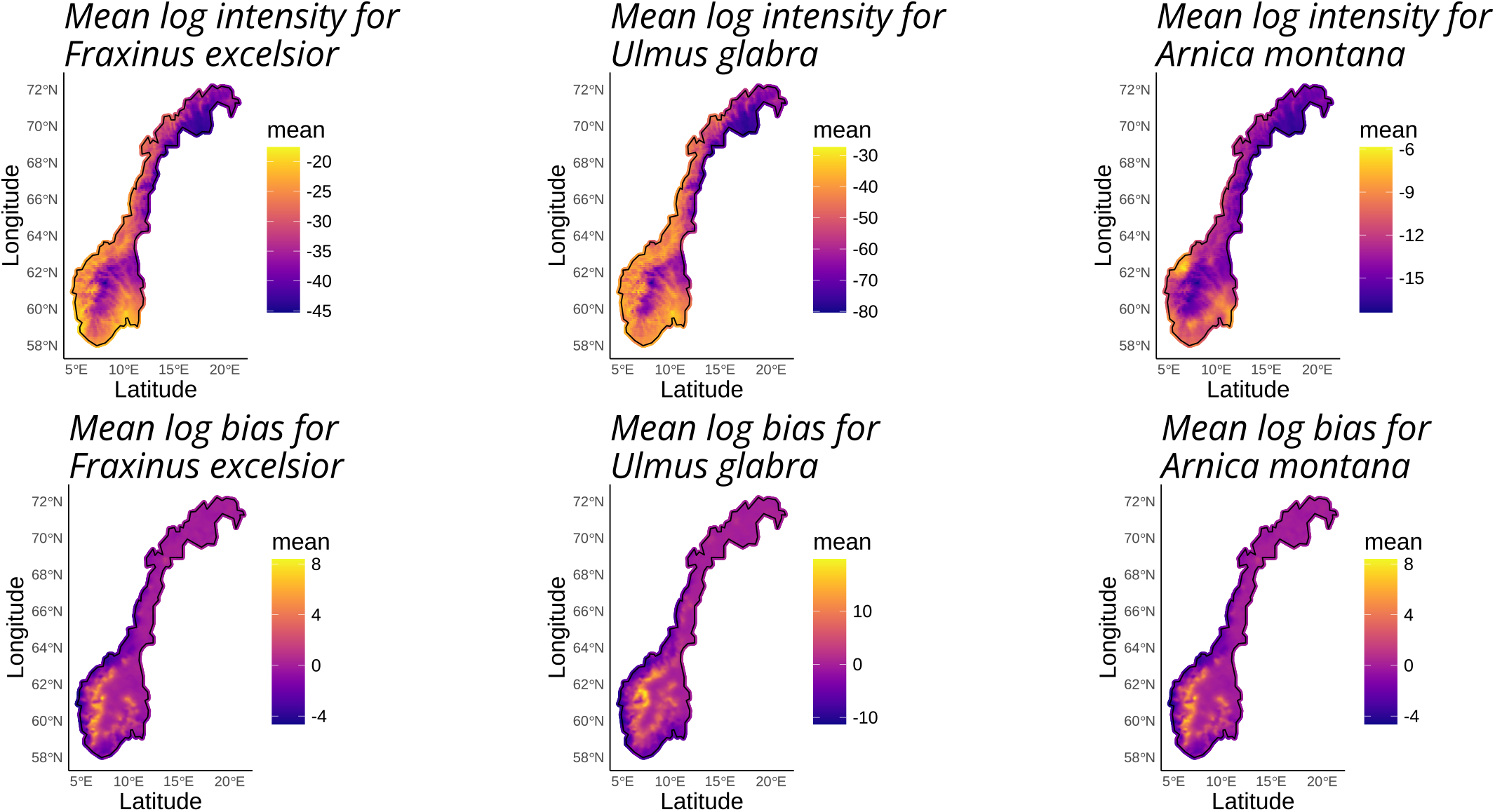
Mean of the log predicted intensity and bias for the three species, A. montana, F. excelsior and U. glabra across Norway, from ISDMs based on presence absence and presence-only data for each respective species, and annual mean temperature.

## 4 Discussion

Reproducible workflows are imperative in ecology to ensure full transparency, generality and consistency in the results of research studies. Even though it is impossible to perfectly replicate observation studies in the field, ecologists need to focus on producing reproducibility in areas of their studies where they can, such as computational reproducibility (Powers & Hampton, 2019). Employing methods such as data integration for SDMs may assist in producing such workflows since they minimize the challenges related to combining observations from heterogeneous study systems.

A package like *intSDM* provides the toolkit to produce a open, reproducible, and flexible workflow integrating different data types from multiple sources to use integrated SDMs to construct maps of species distributions. Specifically, it facilitates the combination of presence-only, presence-absence and counts data, which has the potential to increase the quality of SDMs. A benefit of presenting the workflow through a *CRAN*-submitted *R*-package is that certain forms of software reproducibility such as continuous integration, adequate documentation and unit tests are required to ensure the package’s functions and scripts run properly on a variety of platforms.

In the presented case study, we used the package to produce maps of predicted species distributions for three red listed vascular plant species in Norway. Such a map could provide a knowledge basis for biodiversity conservation. Furthermore, reproducible workflows like *intSDM* have the potential to implement and thus promote best practice standards in the model building process of SDMs (such as those proposed by Araújo *et al*. [2019] and Zurell *et al*. [2020]), which have been sketched out in order to improve the overall transparency of reproducible workflows, raise awareness of model quality and reduce potential misuse of model outputs.

However, future research needs to be considered for *intSDM* in order to make it more flexible for different analyses and easier to use in the future. Incorporating different elements such as adding a temporal component to the model is requisite in order to understand the dynamics of species over time. In addition, creating a graphical user interface (GUI) would make the package more attractive for users with minimal coding skills, and would be able to output visual summaries of each component of our workflow. The extension to this package should improve the upstream components of this workflow: obtaining data from a many different online data repositories (not only GBIF), consider more data types other than presence-only and presence-absence data and being able to create a coherent data structure for each dataset with standardized metadata – even if sampling protocols vary greatly between them.

Despite the substantial added value of integrating several sources of data in SDMs, the use of disparate data may still cause some challenges. Presence-only data are often the most abundant data source, but observations are likely to be biased towards conveniently located, densely populated areas (Sicacha-Parada *et al*., 2021) or certain species (Amano *et al*., 2016; Troudet *et al*., 2017). Integrating presence-absence and presence-only data can increase the quality of SDMs, but how much depends on the amount of available presence-absence data, and the ability to correct for bias in the presence-only data (Simmonds *et al*., 2020). Datasets of different sizes and quality affect results by producing biased estimates towards the more abundant data sources (Zipkin *et al*., 2021). Common solutions involve sub-sampling or down-weighting the more abundant datasets; however these methods are often subjective and can lead to different conclusions (Maunder & Piner, 2017). For species with fewer occurrences, issues connected to low data availability may arise, such as inflating uncertainty of model predictions obtained for these species. Despite this, the modelling framework here could be used as a potential solution. Plotting species intensities against rare species occurrences could be used to show where rare species are likely to be found, but haven’t been sampled yet, providing a good basis for more investigation and fieldwork to expand knowledge of the species distribution.

For an open workflow to be truly reproducible, data need to be open too; therefore there is an ongoing effort towards making biodiversity data available. Data from GBIF are FAIR (Wilkinson *et al*., 2016) and formatted according to prevailing standards such as Darwin Core (Wieczorek *et al*., 2012). Despite this, there was deficient documentation and metadata for species occurrence data in our case study, requiring consultation with data owners to understand the collection methods. A lack of necessary documentation is not unique to this specific data set, highlighting that there are still challenges to resolve to make biodiversity science truly open and reproducible. Understanding the output of the model properly is essential when using a tool made by others. This is not unique for *intSDM* but rather applies to all available *R*-packages or tools for data analyses. Providing properly documented scripts and a vignette illustrating the use of *intSDM* is the first step to mitigate this issue.

It is also important to understand the data included in SDMs in order to account for weaknesses and data quality differences between the datasets, and to make sure data are properly prepared before use. Knowledge about the species or system under study is also key to identify and include the necessary data to produce high quality SDMs, and make sure that the produced output is sensible. The package presented here maximises usefulness of available open source data through integration, and provides a structured framework for other users to follow in order to continue producing open, reproducible workflows.

## Acknowledgements

We are grateful to Anders Finstad and Bob O’Hara for their supervision and input on the manuscript, and Rune Sørås for his contributions to the early stages of the methodology development.

## Data availability

All code and data used in this manuscript are freely available in the *R* package, *intSDM*, which is available on the Comprehensive R Archive Network: https://CRAN.R-project.org/package=intSDM (Mostert *et al*., 2022). The *R* code is also available on GitHub: https://github.com/PhilipMostert/intSDM. The version of the package used for this manuscript (v2.1.0) is archived on Zenodo; https://doi.org/10.5281/zenodo.8430035 (Mostert *et al*., 2024).

## Author contributions

All authors: Conceptualization, Methodology, Validation, Writing - Original Draft, Writing - Review & Editing. Additionally, WK: Visualization, PM: Software.

